# Endocytosis of ALK promotes glucose uptake in *ALK*-amplified neuroblastoma

**DOI:** 10.64898/2026.08.04.742396

**Authors:** Ryouhei Tsutsumi, Shoko Hikage, Shinichi Kiyonari, Ryuichi Sakai

## Abstract

Activated receptor tyrosine kinases (RTKs), such as epidermal growth factor receptor (EGFR) and anaplastic lymphoma kinase (ALK), trigger intracellular signaling while undergoing receptor endocytosis. We recently identified a noncanonical mechanism in which RTK-containing endocytic vesicles deliver extracellular glucose to hexokinases associated with the outer mitochondrial membrane, thereby promoting cellular glucose uptake. Whether this mechanism contributes to cancer metabolism, however, remains unknown. Here, using neuroblastoma cell lines with distinct *ALK* alterations, we investigated the role of ALK endocytosis in glucose uptake. *ALK*-amplified, but not *ALK*-mutant, neuroblastoma cells exhibited a ∼40– 50% reduction in glucose uptake following inhibition of ALK or receptor endocytosis. This process was independent of the ERK MAPK and PI3K–AKT pathways but required dynamin-dependent endocytosis, cytoplasmic dynein, and GLUT1. Overexpressed ALK constitutively co-endocytosed with GLUT1 into vesicles transported to mitochondria. Inhibition of ALK activity or endocytosis suppressed glucose uptake without producing an additive effect, indicating that both function within the same pathway. Furthermore, disruption of the endocytic machinery selectively impaired the growth of *ALK*-amplified neuroblastoma cells. These findings identify ALK endocytosis as a major regulator of glucose uptake in *ALK*-amplified neuroblastoma and suggest that RTK endocytosis represents a previously unrecognized metabolic vulnerability that may be therapeutically exploitable in RTK-driven cancers.

## Introduction

Neuroblastoma is a major extracranial solid tumor of childhood and arises from neural crest-derived progenitors of the sympathetic nervous system, most frequently in the adrenal medulla or paraspinal sympathetic ganglia (1). *MYCN* amplification (20–25% of cases) is the most common genetic alteration associated with aggressive neuroblastoma (1–3), followed by activating point mutations in *ALK* (approximately 10%) and *ALK* amplification (2–5%), the latter frequently co-occurring with *MYCN* amplification (1, 4–8). The *ALK* gene encodes anaplastic lymphoma kinase (ALK), a receptor tyrosine kinase (RTK) (4, 9), which is activated by extracellular ligands, including ALKAL1 and ALKAL2 (10–13).

RTKs bind their cognate growth factors at the plasma membrane and undergo autophosphorylation, thereby initiating intracellular signaling while simultaneously triggering receptor endocytosis (9, 14). Endocytosis results in incorporation of RTKs into endocytic vesicles that subsequently fuse with early endosomes and mature into late endosomes and lysosomes (14). Alternatively, internalized RTKs can be recycled back to the plasma membrane through several pathways, including that involving recycling endosomes (14). Oncogenic activation or overexpression of RTKs, including epidermal growth factor receptor (EGFR), human epidermal growth factor receptor 2 (HER2), and ALK, drives ligand-independent receptor activation and downstream signaling, thereby promoting the highly proliferative phenotype of cancer cells (5, 9, 15). On the other hand, the effects of oncogenic RTK mutations or overexpression on receptor endocytosis vary considerably among RTK family members and depend on the cellular context (16, 17).

We recently reported a noncanonical mechanism by which RTK-containing endocytic vesicles function as carriers that deliver extracellular glucose to glycolytic enzymes located adjacent to mitochondria, thereby enhancing cellular glucose uptake in growth factor-stimulated mouse embryonic fibroblasts (18). Glucose incorporated within endocytic vesicles is transported to sites adjacent to mitochondria and released into the cytoplasm through the co-endocytosed glucose transporter GLUT1 (Fig. S1). This mechanism was observed in response to multiple growth factors, including platelet-derived growth factor (PDGF), basic fibroblast growth factor (bFGF), and hepatocyte growth factor (HGF), and requires RTK activation, RTK/GLUT1 co-endocytosis, and dynein-mediated vesicle transport but is independent of canonical ERK MAPK and PI3K-AKT signaling (18–20), unlike previously described mechanisms that increase glucose uptake by promoting plasma membrane localization or expression of glucose transporters (21–23).

Whether this mechanism also contributes to the metabolic phenotype of RTK-driven cancers has not been investigated. In the present study, we used a panel of neuroblastoma cell lines with distinct *ALK* genetic alterations to determine whether RTK endocytosis-dependent glucose uptake also occurs in cancer cells. Our results demonstrate that a substantial fraction of glucose uptake depends on ALK endocytosis in *ALK*-amplified, but not necessarily *ALK*-mutant, neuroblastoma cells.

## Materials/Subjects and Methods

### Antibodies and chemical compounds

Rabbit polyclonal antibody against the cytoplasmic region (amino acids 1379–1524) of human ALK was generated in the Sakai laboratory as previously described (24). Mouse monoclonal anti-ALK antibody (MAB42101) was purchased from R&D Systems (Minneapolis, MN, USA). Rabbit monoclonal antibodies against GAPDH (D16H11, #5174), AKT (C67E7, #4691), pT308 AKT (D25E6, #13038), pS473AKT (D9E, #4060), ERK1/2 (137F5, #4695), phospho ERK1/2 (D13.14.4E, #4370), and rabbit polyclonal anti-pY1604 ALK antibody (#3341) were purchased from Cell Signaling Technology (Danvers, MA, USA). Rabbit monoclonal anti-GLUT1 (ab115730), anti-dynamin 1 (ab52611), anti-dynamin 3 (ab134925) antibodies, rabbit polyclonal anti-dynamin 2 antibody (ab3457), and Alexa Fluor-conjugated anti-TOMM20 rat monoclonal antibody (ab309167) were from Abcam (Cambridge, UK). Mouse anti-phosphotyrosine monoclonal antibody cocktail 4G10 Platinum (05-1050) was purchased from Sigma-Aldrich (St. Louis, MO, USA). Rabbit polyclonal anti-HA-tag antibody was from MBL. Goat polyclonal anti-EEA1 (sc-6414) antibody was purchased from Santa Cruz Biotechnology (Dallas, TX, USA) but is currently discontinued. Alexa Fluor-conjugated secondary antibodies were purchased from Thermo Fisher Scientific (Waltham, MA, USA).

BAY-876 (6199) was purchased from Tocris Bioscience (Bristol, UK); selumetinib (AZD6244, S1008) from Selleck (Houston, TX, USA); buparlisib (BKM120, HY-70063) from MedChemExpress (Monmouth Junction, NJ, USA); daraxonrasib (AMBH9A9ACD45), ciliobrevin D (250401) and dynarrestin (SML2332) from Sigma-Aldrich.

### Cell culture and transfection

NB1, SK-N-SH, and NB69 were obtained from RIKEN BRC; SH-SY5Y, and SK-N-AS from ATCC, LAN5, and SK-N-FI from the Childhood Cancer Repository (Texas Tech University Health Sciences Center, Lubbock, TX, USA); IMR32 from JCRB (National Institutes of Biomedical Innovation, Health and Nutrition, Japan). NB39nu (from National Cancer Center Research Institute, Japan) (25), TNB1 (from Human Science Research Resource Bank, Japan), and SK-N-MC (from Human Science Research Resource Bank, Japan) were maintained in the Sakai Laboratory. HEK293T cells were obtained from ATCC. All cells were confirmed as mycoplasma-negative by the PCR method as reported previously (26).

Neuroblastoma cells were cultured in RPMI 1640 medium supplemented with 10% fetal bovine serum (FBS), non-essential amino acid solution (Nacalai Tesque, Kyoto, Japan, #06344-56), and Sodium Pyruvate Solution (Nacalai Tesque, #06977-34). HEK293T cells were cultured in Dulbecco-modified Eagle’s medium (DMEM, Nacalai Tesque, #08458-16) supplemented with 10% FBS.

Neuroblastoma cells were transfected with the indicated siRNAs using Lipofectamine RNAiMAX (Thermo Fisher), according to the manufacturer’s protocol. Pre-designed siRNAs targeting human *DNM1* (SASI_Hs01_00189738, SASI_Hs01_00189739), *DNM2* (SASI_Hs01_00227659, SASI_Hs01_00227662), and *DNM3* (SASI_Hs01_00015190, SASI_Hs01_00015192), and a validated negative control siRNA (SIC001) were purchased from Sigma-Aldrich.

### SDS-PAGE and Immunoblotting

Cells were lysed in SDS lysis buffer (50 mM Tris-HCl pH7.5, 100 mM NaCl, 1 mM EDTA, 1% SDS, 10 mM NaF, 10 mM β-glycerophosphate, 2 mM Na_3_VO_4_). Cell lysates were subjected to SDS-PAGE, followed by transfer to Immobilon-P PVDF membranes (Millipore, Burlington, MA, USA). Membranes were blocked in 1% BSA/TBS containing 0.1% Tween20 for 30 min, and treated with primary antibodies in blocking buffer overnight at 4°C, followed by treatment with HRP-conjugated secondary antibodies (Cell Signaling Technology, #7074, #7076) for 1 h. Bands were visualized using Chemi-Lumi One Super (Nacalai Tesque, #02230-30) according to the manufacturer’s protocol, and images were obtained using an Amersham™ ImageQuant™ 800 (Cytiva, Marlborough, MA, USA) as unsaturated 16-bit TIFF images.

### Glucose uptake assays

Glucose uptake was quantified using the Glucose Uptake-Glo™ Assay kit (Promega, Madison, WI, USA) according to the manufacturer’s protocol with some modifications. For inhibitor treatment, neuroblastoma cells (1 x 10^4^ per well in a 96-well plate) were seeded and cultured for 48 h, followed by inhibitor or conditioned medium treatments by changing medium to DMEM containing 0.1% DMSO with or without indicated concentrations of inhibitors for 10 min. Cells were then rinsed twice with PBS, and incubated in PBS containing 1 mM 2DG at 37°C for 10 min. Vehicle or inhibitor was also added to the solutions for rinsing and 2DG treatment. For siRNA experiments, cells were reverse-transfected at the time of seeding with the indicated siRNAs using Lipofectamine RNAiMAX (Thermo Fisher), and cultured for 48 h, followed by glucose uptake assay. Cellular glucose uptake was terminated by adding acidic Stop Buffer, followed by addition of Neutralization Buffer. The lysates were mixed with the 2DG6P detection reagent, containing G6P dehydrogenase, NAD+, reductase, proluciferin, ATP and recombinant luciferase, and incubated at RT for 1 h. Luminescence signals were quantified using a SpectraMaxiD5 plate reader (Molecular Devices). Absolute amounts of cellular 2DG6P formation were calculated based on signals of 2DG6P standards, setting the signals of 2DG-untreated cells as the background. 2DG6P formation was normalized by amounts of protein per well.

### Conditioned medium

Human *ALKAL2* coding sequence with a 5’-HA-tag sequence was commercially synthesized and cloned into pcDNA3.1 vector. HEK293T cells (2.5 x 10^6^ per dish) were seeded on 10-cm dishes in DMEM containing 10% FBS, and transfected with an empty vector or HA-tagged ALKAL2 expression vector using Lipofectamine 3000 (Thermo Fisher), according to the manufacturer’s protocol. At 6 h after transfection, the medium was replaced with FBS-free DMEM, and cells were cultured for 40 h. The conditioned medium was then centrifuged 800 x g for 5 min to remove cells and debris.

### Immunofluorescence

Cells (5 x 10^4^) were seeded on 12 mm, poly-D-lysine-or atelocollagen acidic solution (Koken, Tokyo, Japan, I-PC)-coated circular glass coverslips. Cells were rinsed with PBS and fixed in 4% paraformaldehyde/PBS for 10 min at RT, permeabilized with 0.1% Triton X-100, 0.2% BSA/PBS for 10 min, and blocked with 1% BSA/PBS for 30 min, followed by sequential primary antibody and Alexa fluor-conjugated secondary antibody (and Alexa fluor-conjugated anti-TOMM20 antibody) treatments as indicated for 1 h each. Coverslips were then mounted on glass slides using ProLong Gold containing DAPI (Thermo Fisher). Images were obtained using an LSM980 confocal microscopy system (Zeiss, Oberkochen, Germany) as unsaturated 16-bit TIFF images. Confocal microscopes were set to obtain 1 Airy Unit (AU), and single-plane images are presented in figures. Image processing, including pseudo-coloring, was performed using FIJI/ImageJ Ver. 1.54r (27).

### Cell growth assay

Neuroblastoma cells (5 x 10^4^ per well) were seeded on 48-well plates in RPMI 1640 medium containing 10% FBS for 2 days, followed by transfection using Lipofectamine RNAiMAX (Thermo Fisher), according to the manufacturer’s protocol. Cells were then incubated at 37°C in a CO_2_ incubator for 5 days. Medium was changed at 48, 72, and 96 h after transfection. Cells were rinsed with PBS, followed by staining with 0.5% crystal violet solution containing 25% methanol for 30 min. Culture plates were rinsed with tap water and air dried. Plates were then scanned as 16-bit TIFF images.

### Statistics and reproducibility

No statistical method was used to predetermine sample sizes. Samples were not randomized. The investigators were not blinded to allocation during experiments or outcome assessment. Sample sizes and statistical tests for each experiment are denoted in the figure or legends. Each experiment was performed at least twice per condition of the experiment, and representative images from one of the biological replicates are shown in each panel.

Statistical analysis was performed by using paired two-tailed t-tests (Figs. 1B, 1C, 1D, 4B), repeated measures one-way ANOVA with the Geisser-Greenhouse correction, and Dunnet’s multiple comparisons test with individual variances computed for each comparison (1E, 4A), or Sidak’s multiple comparisons test with individual variances computed for each comparison (4C, 4D) by using GraphPad Prism 8 Ver. 8.4.3 (GraphPad Software, Boston, MA, USA), where appropriate. Precise P values can be found in the figures.

**Fig. 1.**
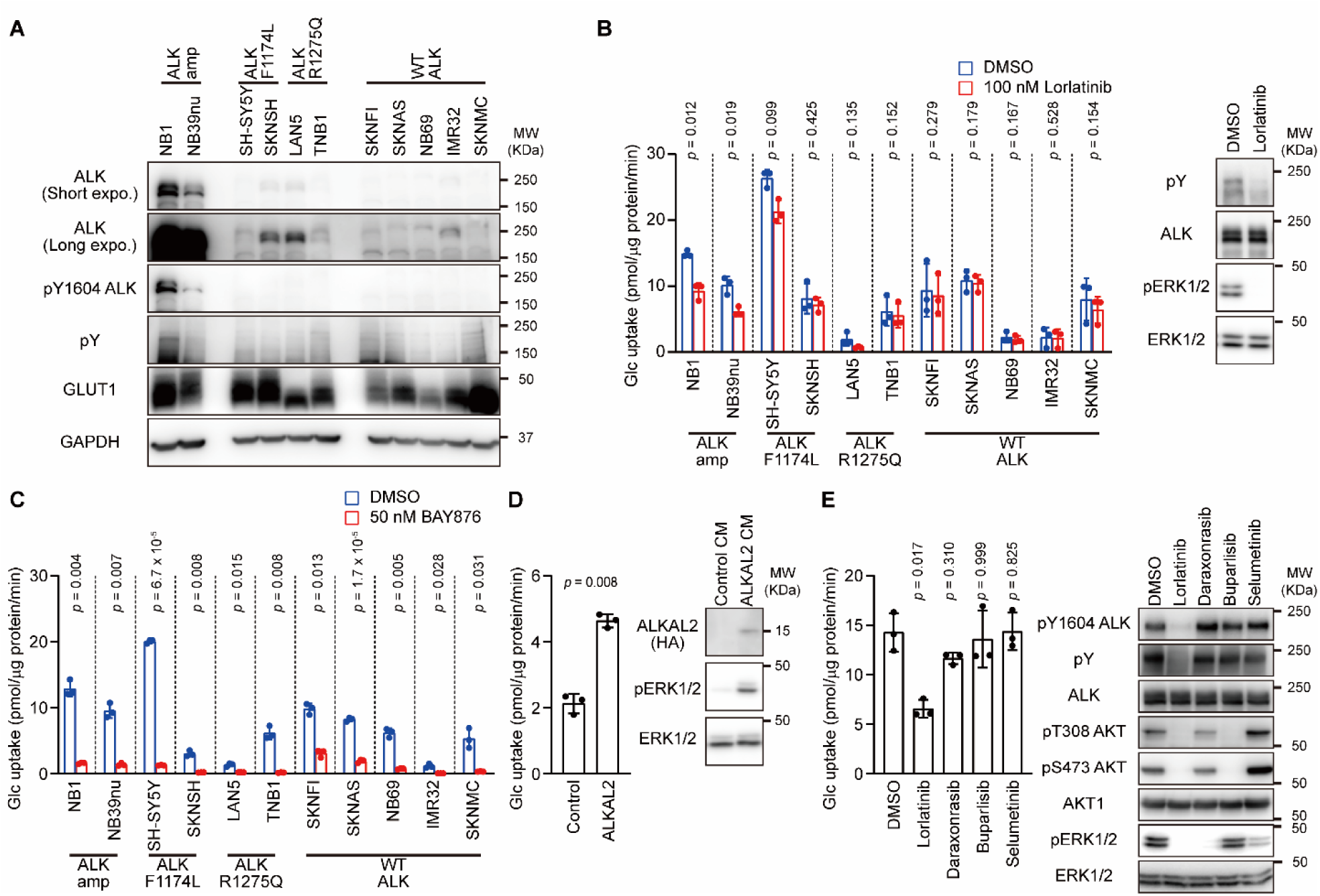
ALK-and GLUT1-dependent glucose uptake in ALK-amplified neuroblastoma cell lines. **A** Lysates from neuroblastoma cell lines were subjected to immunoblotting with indicated antibodies. Representative images are shown from one of 2 independent experiments. **B** Cells were pre-treated with or without ALK-specific inhibitor lorlatinib (1 μM) for 10 min, and then subjected to glucose uptake assay in the presence of 2DG with or without lorlatinib for 10 min. Graph shows 2DG6P normalized to total cellular proteins (left). Bars and error bars in the graph show means of 3 independent biological replicates and ± SD (biological replicates, n = 3) for each cell line. P values were calculated using paired two-tailed t tests. Lysates from NB1 cells treated with or without lorlatinib for 10 min were subjected to immunoblotting with indicated antibodies (right). **C** Cells were pre-treated with or without GLUT1-specific inhibitor BAY-876 (50 nM) for 10 min, and then subjected to glucose uptake assay in the presence of 2DG with or without BAY-876 for 10 min. Graph shows 2DG6P normalized to total cellular proteins. Bars and error bars in the graph show means of 3 independent biological replicates and ± SD (biological replicates, n = 3) for each cell line. P values were calculated using paired two-tailed t tests. **D** IMR32 cells were pre-treated with conditioned medium of HEK293T cells transfected with HA-tagged ALKAL2-expressing vector (ALKAL2 CM) or control expression vector (control CM) for 10 min, and then subjected to glucose uptake assay in the presence of 2DG. Graph shows 2DG6P normalized to total cellular proteins (left). Bars and error bars in the graph show means of 3 independent biological replicates and ± SD (biological replicates, n = 3). P values were calculated using paired two-tailed t test. Lysates were subjected to immunoblotting with indicated antibodies (right). **E** NB1 cells were pre-treated with lorlatinib (1 μM), daraxonrasib (1 μM), buparlisib (1 μM), selumetinib (1 μM), or vehicle for 10 min, and then subjected to glucose uptake assay in the presence of 2DG for 10 min. Graph shows 2DG6P normalized to total cellular proteins (left). Bars and error bars in the graph show means of 3 independent biological replicates and ± SD (biological replicates, n = 3) for each cell line. P values were calculated using repeated measures one-way ANOVA with the Geisser-Greenhouse correction, and Dunnet’s multiple comparisons test with individual variances computed for each comparison. Lysates were subjected to immunoblotting with indicated antibodies (right).

### Large Language Models (LLMs)

Portions of the manuscript were edited for English language and clarity using ChatGPT (OpenAI, GPT-5.5), with all scientific content reviewed and verified by the authors.

## Results

### Constitutively activated ALK in ALK-amplified neuroblastoma cell lines

To investigate the role of ALK and its endocytosis in glucose uptake by neuroblastoma, we analyzed a panel of 11 neuroblastoma cell lines harboring distinct *ALK* genetic alterations, including *ALK* amplification (NB1 and NB39nu), the *ALK F1174L* mutation (SH-SY5Y and SK-N-SH), the *ALK R1275Q* mutation (LAN5 and TNB1), or wild-type *ALK* (SK-N-FI, SK-N-AS, NB69, IMR32, and SK-N-MC). Among these, NB1, NB39nu, LAN5, TNB1, and IMR32 harbor *MYCN* amplification. The majority of activating ALK mutations in neuroblastoma occur within the activation loop of the kinase domain at residues F1174, F1245, and R1275 (4). The F1174L and the R1275Q mutations are well-characterized activating mutations (5, 28).

Immunoblotting of equal amounts of total cellular protein demonstrated markedly abundant ALK expression in the *ALK*-amplified cell lines NB1 and NB39nu, as expected, whereas several cell lines harboring mutant or wild-type *ALK*, including SK-N-SH, LAN5, and IMR32, exhibited relatively low but detectable levels of ALK (Fig. 1A). Immunoblotting with an antibody recognizing ALK phosphorylated at Y1604, one of the autophosphorylation sites that reflects ALK activation, as well as with the 4G10 anti-phosphotyrosine (pY) antibody, revealed ALK autophosphorylation in NB1 and NB39nu cells, indicating constitutive ligand-independent ALK activation, at least in these cells (Fig. 1A). ALK autophosphorylation was barely detectable in the remaining cell lines, including those harboring mutant *ALK,* when compared with *ALK*-amplified cells. Meanwhile, expression levels of GLUT1 vary among cell lines, with no obvious correlation with ALK expression or *MYCN* amplification.

### ALK-and GLUT1-dependent enhanced glucose uptake in neuroblastoma cell lines

We next examined whether acute inhibition of ALK affects glucose uptake in neuroblastoma cells by using the third-generation ALK inhibitor lorlatinib, which potently inhibits both wild-type ALK and the majority of oncogenic ALK mutants (29, 30). Following a 10-min pre-treatment with lorlatinib, cells were incubated with the glucose analog 2-deoxyglucose (2DG), which is transported into cells by glucose transporters and phosphorylated by hexokinases but is not further metabolized through glycolysis. Cellular glucose uptake was quantified by measuring the amounts of phosphorylated 2DG (2DG6P). Pre-treatment with lorlatinib (1 μM) for 10 min significantly reduced glucose uptake during the subsequent 10 min in NB1 and NB39nu by approximately 40%, whereas no significant reduction was observed in most cell lines harboring wild-type or mutant *ALK* (Fig. 1B). SH-SY5Y and LAN5 exhibited a modest decrease in glucose uptake by lorlatinib, but these changes did not reach statistical significance (Fig. 1B). Immunoblotting confirmed that treatment with 1 μM lorlatinib for 10 min efficiently suppressed ALK autophosphorylation in NB1 cells (Fig. 1B). Although lorlatinib is also known to inhibit ROS1, publicly available databases, such as The Human Protein Atlas, indicate that neuroblastoma cell lines generally express substantially lower mRNA levels of *ROS1* than *ALK*. Thus, approximately 40% of glucose uptake in *ALK*-amplified neuroblastoma cells depends on ALK activity. Notably, this fraction is unlikely to result from ALK-dependent transcriptional regulation and de novo protein synthesis because of the rapid effect. Glucose uptake in neuroblastoma cells is most likely mediated by GLUT1, as treatment with 50 nM BAY-876, which selectively inhibits GLUT1 among the major glucose transporters (GLUT1–4) at this concentration (31), blocked the formation of 2DG6P by more than 90% in all cell lines except SK-N-FI and SK-N-AS (Fig. 1C). Thus, the ALK-dependent glucose uptake in *ALK*-amplified neuroblastoma cells is mediated primarily through GLUT1.

To explore the possibility that ALK activation enhances cellular glucose uptake, we next investigated whether activation of wild-type ALK is sufficient to stimulate glucose uptake. IMR32 cells, which express wild-type ALK (Fig. 1A), were treated with conditioned medium collected from HEK293T cells transiently transfected with an ALKAL2 expression vector or an empty control vector. Activation of ALK by ALKAL2 was confirmed by increased ERK1/2 phosphorylation (Fig. 1D). Pre-treatment with ALKAL2-conditioned medium for 10 min increased glucose uptake approximately twofold in IMR32 cells compared with control conditioned medium (Fig. 1D).

Collectively, these results demonstrate that constitutive ALK activation enhances glucose uptake, particularly in *ALK*-amplified, but not necessarily *ALK*-mutant, neuroblastoma cells. Glucose uptake in *ALK*-amplified cells consists of an ALK-dependent component, accounting for approximately 40% of total glucose uptake, and an ALK-independent basal component, both of which are mediated primarily by GLUT1. To explore the underlying mechanism of ALK-dependent glucose uptake, we used NB1 as a representative model of *ALK*-amplified neuroblastoma cells in subsequent experiments.

### ALK-dependent enhanced glucose uptake does not require the ERK MAPK or PI3K-AKT pathways

Activation of ALK, like that of other RTKs, induces autophosphorylation of cytoplasmic tyrosine residues, leading to recruitment of multiple signaling proteins and activation of downstream pathways, including the RAS-ERK MAPK and PI3K-AKT pathways. However, treatment of NB1 cells with the RAS(ON)-specific inhibitor daraxonrasib, the MEK1/2 inhibitor selumetinib, or the pan-PI3K inhibitor buparlisib had only a marginal suppression of glucose uptake in contrast to lorlatinib (Fig. 1E). Immunoblotting confirmed effective inhibition of the intended signaling pathways, as demonstrated by reduced phosphorylation of ERK1/2 and AKT following treatment with these inhibitors (Fig. 1E). These observations indicate that the ALK-dependent component of glucose uptake in *ALK*-amplified neuroblastoma cells is largely independent of the canonical ERK MAPK and PI3K-AKT pathways.

### Overexpressed ALK undergoes co-endocytosis with GLUT1

Because many RTKs undergo ligand-induced endocytosis (14), we examined whether activation of wild-type ALK similarly promotes receptor internalization. IMR32 cells were stimulated with ALKAL2-conditioned medium and immunostained for ALK and the early endosome marker EEA1. Following ligand stimulation, ALK accumulated in intracellular vesicular structures that colocalized with EEA1, demonstrating that activated ALK undergoes endocytosis through the canonical early endosomal pathway (Fig. 2A). We next investigated the subcellular localization of ALK in *ALK*-amplified NB1 cells. ALK was detected in cytoplasmic vesicles that colocalized with EEA1 (Fig. 2B), indicating that overexpressed ALK constitutively undergoes endocytosis in these cells.

**Fig. 2.**
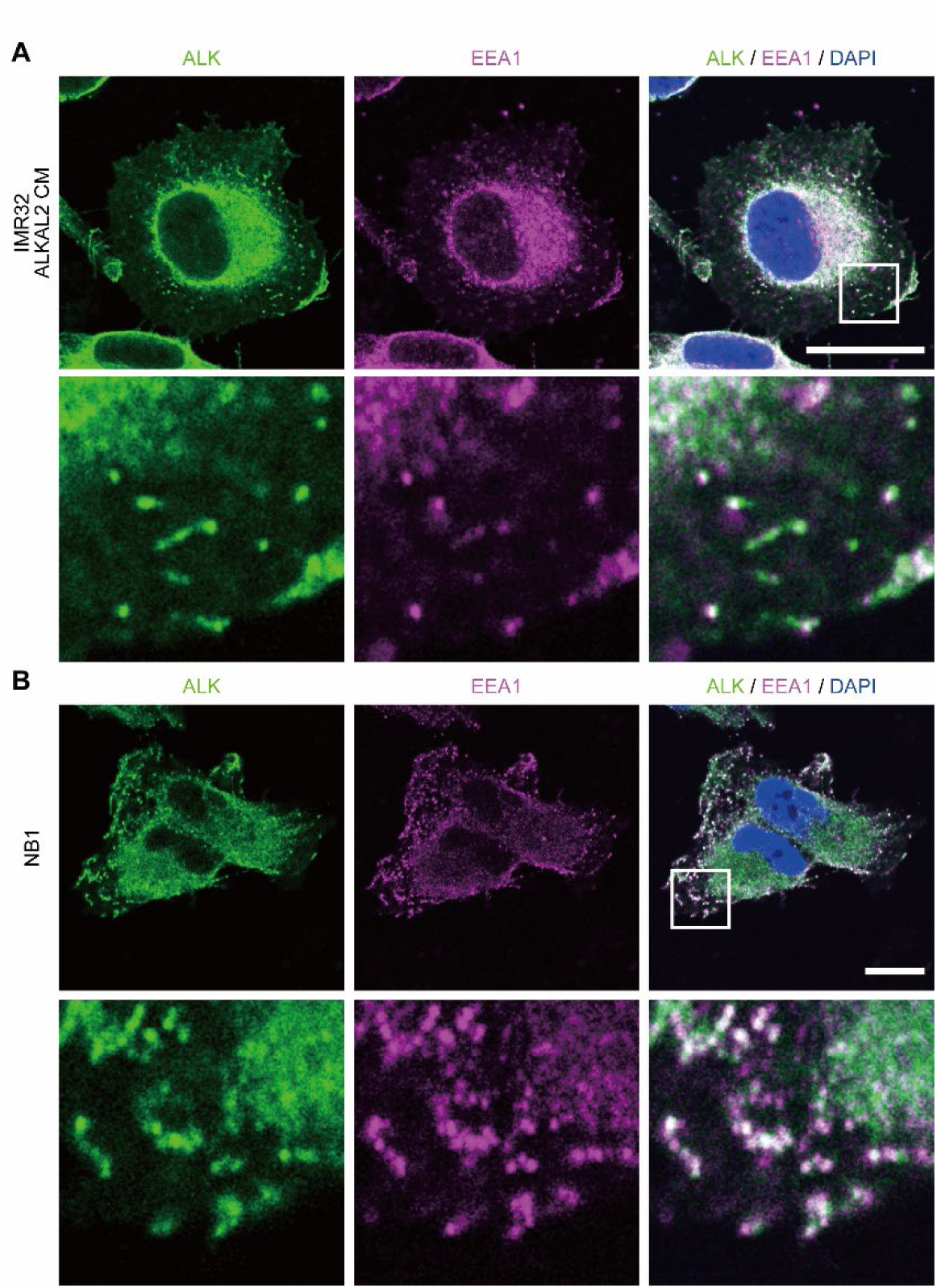
Endocytosis of ALK in neuroblastoma cells. **A** IMR32 cells were pre-treated with conditioned medium of HEK293T cells transfected with ALKAL2-expressing vector (ALKAL2 CM) or control expression vector (control CM) for 10 min, and then subjected to immunostaining with anti-ALK (green) and anti-EEA1 (magenta) antibodies. Nuclei were stained with DAPI (blue). Higher magnification images of the boxed regions are shown on the bottom. B NB1 cells were subjected to immunostaining with anti-ALK (green) and anti-EEA1 (magenta) antibodies. Nuclei were stained with DAPI (blue). Higher magnification images of the boxed regions are shown on the bottom. Scale bars: 10 μm.

These observations suggested that ALK-dependent glucose uptake in *ALK*-amplified neuroblastoma cells may occur through the same mechanism of RTK endocytosis-dependent glucose uptake that we previously described in growth factor-stimulated fibroblasts (18). In that model, activated RTKs and GLUT1 are co-endocytosed with extracellular glucose into endocytic vesicles that are transported to the vicinity of mitochondria. To determine whether ALK-containing vesicles share these characteristics, NB1 cells were co-stained for ALK and GLUT1. As expected, ALK-containing vesicles were extensively colocalized with GLUT1, indicating co-endocytosis of ALK and GLUT1 in *ALK*-amplified neuroblastoma cells (Fig. 3A). Moreover, these vesicles were frequently observed adjacent to mitochondria, visualized by co-staining for the outer mitochondrial membrane protein TOMM20, whereas treatment with lorlatinib markedly reduced the abundance of intracellular ALK-containing vesicles and their localization near mitochondria (Fig. 3B). Together, these findings indicate that constitutively internalized ALK forms GLUT1-positive endocytic vesicles that are transported to mitochondria-proximal regions, supporting the hypothesis that these vesicles mediate the ALK-dependent component of glucose uptake in *ALK*-amplified neuroblastoma cells.

**Fig. 3.**
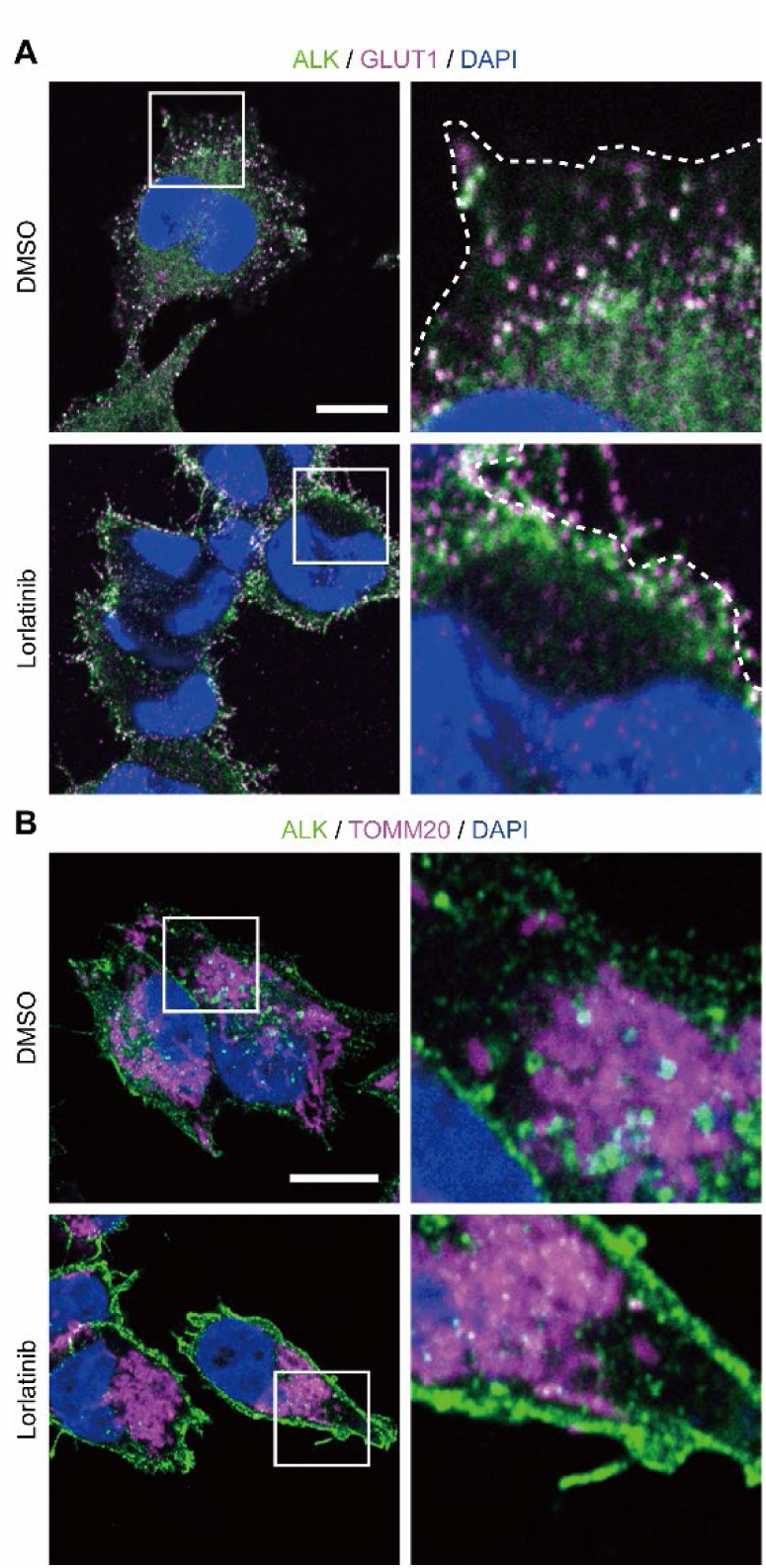
ALK/GLUT1-containing endocytic vesicles in *ALK*-amplified neuroblastoma cells. **A** NB1 cells were subjected to immunostaining with anti-ALK (green) and anti-GLUT1 (magenta) antibodies. Nuclei were stained with DAPI (blue). Higher magnification images of the boxed regions are shown on the right. Dashed lines indicate cell shapes. Scale bars: 10 μm. B NB1 cells were subjected to immunostaining with anti-ALK (green) and anti-TOMM20 (magenta) antibodies. Nuclei were stained with DAPI (blue). Higher magnification images of the boxed regions are shown on the right. Scale bars: 10 μm.

### Endocytosis-dependent glucose uptake in ALK-overexpressing neuroblastoma cells

To determine whether the ALK-dependent glucose uptake requires receptor endocytosis, neuroblastoma cells were treated with siRNAs targeting dynamins, GTPases that mediate the scission of the invaginated plasma membrane during endocytic vesicle formation and are therefore essential for RTK (and ALK) endocytosis (14, 32). In humans, three dynamin isoforms are encoded by *DNM1*, *DNM2*, and *DNM3*. Dynamin 1 and dynamin 3 are predominantly expressed in neuronal tissues, whereas dynamin 2 is ubiquitously expressed. Immunoblot analysis confirmed the expression of all three dynamin isoforms in NB1 cells and efficient knockdown by two independent siRNAs targeting each isoform (Fig. 4A). Silencing either *DNM1* or *DNM2* significantly reduced glucose uptake by approximately 40–50%, whereas knockdown of *DNM3* had no effect (Fig. 4A). Thus, both dynamin 1 and dynamin 2 are required for efficient glucose uptake in NB1 cells. The magnitude of inhibition was comparable to that produced by ALK inhibition with lorlatinib (Figs. 1B and 1E). Simultaneous knockdown of all three dynamin isoforms did not further reduce glucose uptake beyond that observed following depletion of either dynamin 1 or dynamin 2 alone. This observation suggests either that dynamin 1 and dynamin 2 play nonredundant essential roles in this process, or that overall dynamin expression must exceed a critical threshold to support receptor endocytosis and glucose uptake.

**Fig. 4.**
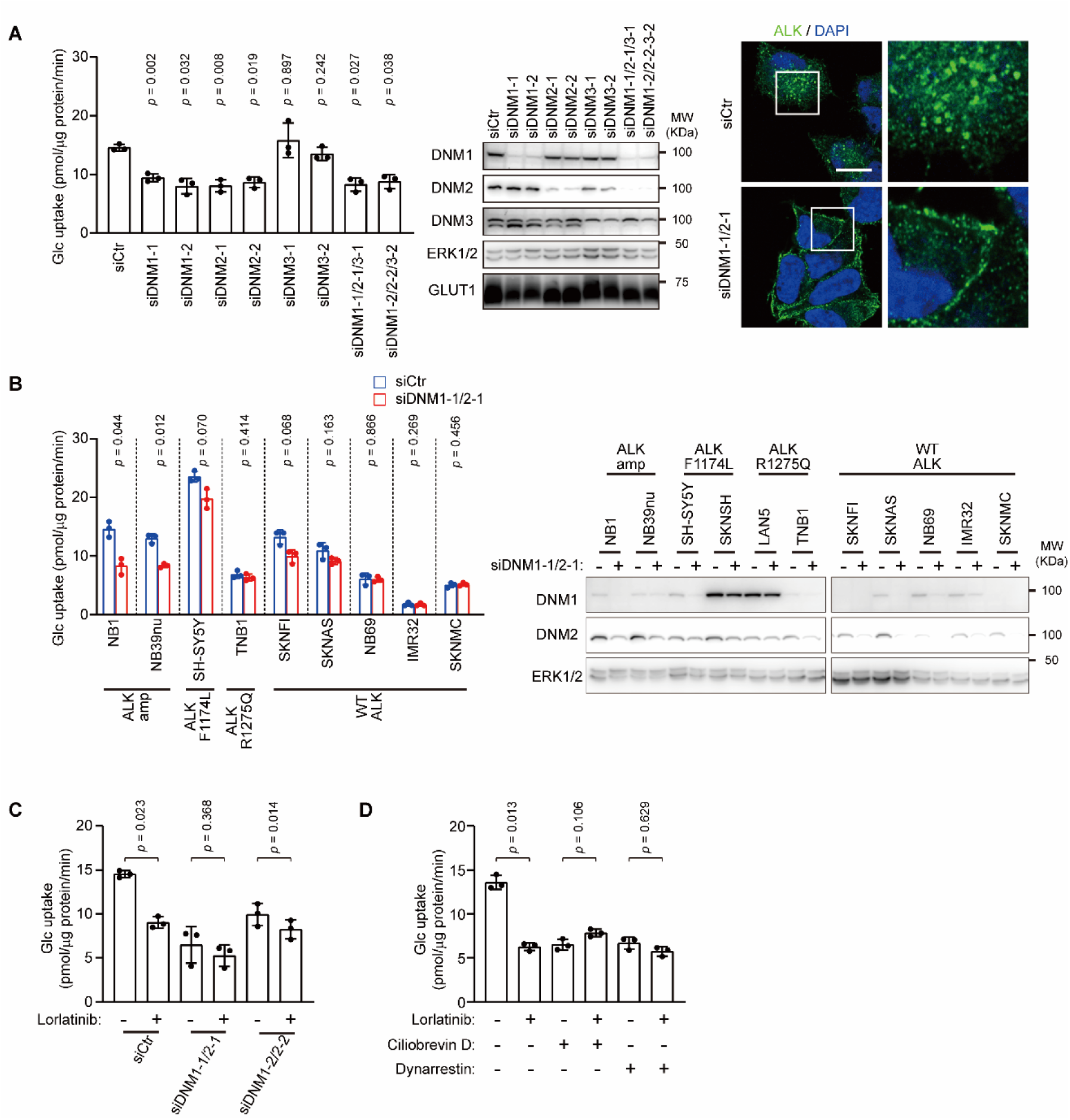
ALK endocytosis-dependent glucose uptake in *ALK*-amplified neuroblastoma cells. **A** NB1 cells transfected with indicated siRNA were subjected to glucose uptake assay in the presence of 2DG. Graph shows 2DG6P normalized to total cellular proteins (left). Bars and error bars in the graph show means of 3 independent biological replicates and ± SD (biological replicates, n = 3). P values were calculated using repeated measures one-way ANOVA with the Geisser-Greenhouse correction, and Dunnet’s multiple comparisons test with individual variances computed for each comparison. Lysates were subjected to immunoblotting with indicated antibodies (middle). NB1 cells transfected with indicated siRNA were immunostained with an anti-ALK antibody (green). Nuclei were stained with DAPI (blue). Higher magnification images of the boxed regions are shown on the right (right). Scale bars: 10 μm. B Neuroblastoma cells transfected with indicated siRNA were subjected to glucose uptake assay in the presence of 2DG. Graph shows 2DG6P normalized to total cellular proteins (left). Bars and error bars in the graph show means of 3 independent biological replicates and ± SD (biological replicates, n = 3) for each cell line. P values were calculated using paired two-tailed t tests. Lysates were subjected to immunoblotting with indicated antibodies (right). **C** NB1 cells transfected with indicated siRNA were pre-treated with or without lorlatinib (1 μM) for 10 min, and then subjected to glucose uptake assay in the presence of 2DG with or without lorlatinib for 10 min. Graph shows 2DG6P normalized to total cellular proteins. Bars and error bars in the graph show means of 3 independent biological replicates and ± SD (biological replicates, n = 3) for each cell line. P values were calculated using repeated measures one-way ANOVA with the Geisser-Greenhouse correction, and Sidak’s multiple comparisons test with individual variances computed for each comparison. **D** NB1 cells were pre-treated with indicated combination of lorlatinib (1 μM), ciliobrevin D (30 μM), and dynarrestin (30 μM) for 10 min, and then subjected to glucose uptake assay in the presence of 2DG for 10 min. Graph shows 2DG6P normalized to total cellular proteins. Bars and error bars in the graph show means of 3 independent biological replicates and ± SD (biological replicates, n = 3) for each cell line. P values were calculated using repeated measures one-way ANOVA with the Geisser-Greenhouse correction, and Sidak’s multiple comparisons test with individual variances computed for each comparison.

We next examined the effect of simultaneous knockdown of dynamin 1 and dynamin 2 in the neuroblastoma cell line panel. SK-N-SH and LAN5 cells were excluded from siRNA experiments hereafter because of their low transfection efficiencies, while efficient knockdown of dynamin 1 and dynamin 2 was confirmed in the remaining cell lines (Fig. 4B). Glucose uptake assays showed that inhibition of endocytosis by silencing dynamins significantly reduced glucose uptake in the *ALK*-amplified cell lines NB1 and NB39nu and produced a modest, although not statistically significant, reduction in SH-SY5Y and SK-N-FI cells (Fig. 4B). In contrast, glucose uptake in the remaining cell lines was largely unaffected by dynamin depletion (Fig. 4B). Notably, the pattern of sensitivity to dynamin depletion closely mirrored that observed following ALK inhibition with lorlatinib (see Fig. 1B), further supporting the conclusion that ALK activity and receptor endocytosis function within the same pathway. Importantly, glucose uptake in most cell lines lacking ALK overexpression was unaffected by dynamin depletion (Fig. 4B), indicating that dynamins are not required for the lorlatinib-insensitive component of glucose uptake in these neuroblastoma cell lines. Furthermore, combined inhibition of ALK activity and dynamin-mediated receptor endocytosis did not further reduce glucose uptake in NB1 cells compared with either intervention alone (Fig. 4C), indicating that ALK kinase activity and receptor endocytosis regulate the same glucose uptake pathway rather than acting through parallel mechanisms. We next examined the role of intracellular vesicle transport. Inhibition of cytoplasmic dynein with either ciliobrevin D or dynarrestin (33) significantly reduced glucose uptake in NB1 cells (Fig. 4D). Simultaneous treatment with lorlatinib produced no additional inhibitory effect, demonstrating that dynein-dependent vesicle transport also functions within the same pathway.

Collectively, these findings demonstrate that ALK-dependent glucose uptake requires dynamin-dependent receptor endocytosis followed by cytoplasmic dynein-mediated vesicle transport. Together, these processes account for approximately 40–50% of total glucose uptake in *ALK*-amplified neuroblastoma cells.

### Survival of ALK-overexpressing neuroblastoma cells requires an intact endocytosis machinery

Finally, we investigated the biological significance of receptor endocytosis in neuroblastoma cell growth. Neuroblastoma cell lines were transfected with control siRNA or siRNAs targeting *DNM1* and *DNM2*, cultured for 5 days, and stained with crystal violet to assess cell growth and survival. Simultaneous depletion of dynamin 1 and dynamin 2 markedly impaired the growth of the *ALK*-amplified cell lines NB1 and NB39nu (Fig. 5). In contrast, little or no effect was observed in neuroblastoma cell lines harboring wild-type or mutant *ALK* (Fig. 5). This pattern closely resembled that observed in the glucose uptake assays (Fig. 4B), in which only *ALK*-amplified cell lines exhibited a strong dependence on dynamin-mediated endocytosis. These findings indicate that an intact endocytic machinery is selectively required for the proliferation and/or survival of *ALK*-amplified neuroblastoma cells, supporting the conclusion that ALK-dependent endocytosis contributes not only to glucose uptake but also to the maintenance of the malignant phenotype.

**Fig. 5.**
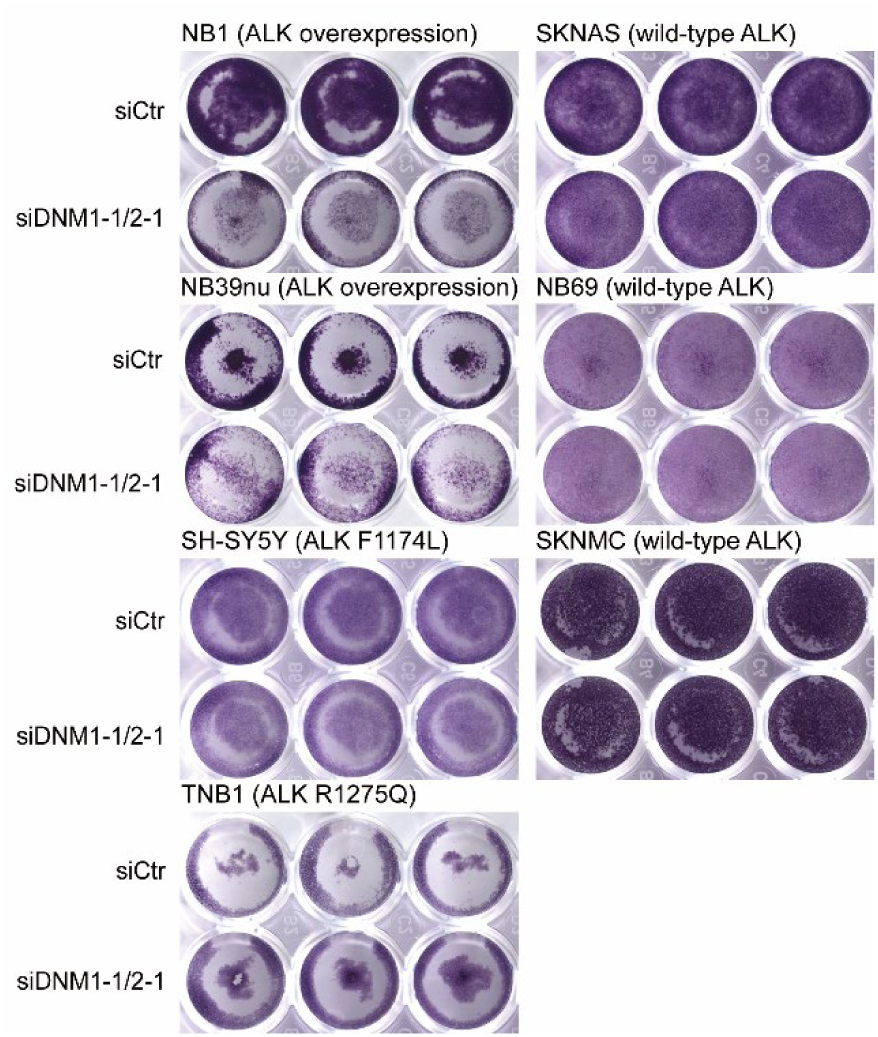
ALK/GLUT1-containing endocytic vesicles in *ALK*-amplified neuroblastoma cells. Neuroblastoma cells transfected with indicated siRNA were cultured for 5 days, and surviving cells were visualized by staining with crystal violet.

## Discussion

Like many adult solid tumors, neuroblastoma exhibits high glucose uptake as detected by positron emission tomography (PET) (34) and displays the Warburg phenotype, characterized by a greater reliance on glycolysis than on mitochondrial oxidative phosphorylation (OXPHOS) (35). This metabolic phenotype has been attributed, at least in part, to *MYCN* overexpression and loss of tumor suppressors such as *TP53*, which enhance the expression of glucose transporters and glycolytic enzymes (35). Consistent with this notion, elevated GLUT1 expression is associated with poor clinical outcome in neuroblastoma, and inhibition of glycolysis has therefore been proposed as a potential therapeutic strategy (36). However, whether ALK overexpression or activating ALK mutations contribute directly to glycolytic regulation in neuroblastoma had remained largely unexplored.

In the present study, we demonstrate that approximately 40–50% of glucose uptake in *ALK*-amplified neuroblastoma cells depends on both ALK kinase activity and an intact endocytic machinery. In contrast, this mechanism was not evident in most neuroblastoma cell lines expressing wild-type or mutant ALK, and no clear association was observed with *MYCN* amplification. Unlike previously described mechanisms that accelerate glycolysis by inducing GLUT4 translocation (21), or transcriptional upregulation of glycolytic genes (22, 23), the present mechanism regulates glucose uptake through receptor endocytosis and therefore operates independently of transcriptional responses.

Several lines of evidence support our conclusion that ALK-containing endocytic vesicles function as carriers for extracellular glucose. First, inhibition of ALK rapidly suppressed glucose uptake within 10 min, making it unlikely that the observed effect results from transcriptional or translational regulation of glucose transporters or glycolytic enzymes. Second, ALK-dependent glucose uptake shared the defining properties of the RTK endocytosis-dependent mechanism previously described in fibroblasts. Finally, inhibition of either ALK kinase activity or receptor endocytosis markedly reduced glucose uptake, whereas simultaneous inhibition produced no additional effect, indicating that both perturbations act within the same pathway. Together with the observed co-endocytosis of ALK and GLUT1, these findings strongly support a model in which ALK-containing endocytic vesicles mediate a substantial fraction of glucose uptake in *ALK*-amplified neuroblastoma cells. Nevertheless, approximately half of total glucose uptake remained resistant to ALK inhibition or suppression of endocytosis, indicating that these cells also maintain a conventional GLUT1-dependent glucose uptake pathway at the plasma membrane.

Our results unexpectedly demonstrated that oncogenic activating mutations of ALK, at least F1174L and R1275Q, did not confer a detectable dependence on ALK-mediated glucose uptake despite their well-established oncogenic kinase activity. We propose that RTK endocytosis-dependent glucose uptake requires a sufficiently large pool of RTK-containing endocytic vesicles and therefore depends not only on receptor activity but also on receptor abundance. Consistent with this model, ALK expression levels in *ALK*-mutant cell lines were considerably lower than those in *ALK*-amplified cells. Previous studies have also shown that activating ALK mutations reduce binding to flotillin-1, thereby attenuating endocytosis-dependent receptor degradation (17). We also cannot exclude the possibility that activating ALK mutations promote glycolytic metabolism through alternative mechanisms, such as transcriptional or translational regulation of metabolic genes.

Our findings may have broader implications beyond neuroblastoma. Amplification or overexpression of RTKs is common in many human cancers, including HER2-positive breast cancer, EGFR-driven cancers, and MET-amplified gastric and hepatocellular carcinomas (37–39). If these receptors similarly utilize endocytic vesicles to facilitate glucose uptake, RTK endocytosis may represent a previously unrecognized metabolic vulnerability shared across multiple RTK-driven malignancies. Therapeutic strategies targeting receptor endocytosis, either alone or in combination with RTK kinase inhibitors, may therefore complement existing approaches that focus primarily on kinase activity and downstream signaling pathways.

Although this study was limited by the relatively small number of available neuroblastoma cell lines, our findings identify receptor endocytosis as an important regulator of glucose metabolism in *ALK*-amplified neuroblastoma and provide evidence that receptor trafficking contributes directly to the metabolic phenotype of RTK-driven cancers. These results expand the functional significance of RTK endocytosis beyond signal transduction and receptor turnover and establish a conceptual framework for investigating endocytosis-dependent metabolic regulation in other oncogene-driven cancers.

## Acknowledgements

This work was supported by Japan Society for the Promotion of Science (JSPS) 24K10317 (to R.T.). R.T. was also supported by the Sumitomo Foundation and the Uehara Memorial Foundation.

## Author Contributions

RT conceptualized and designed research. RT, SH, SK performed research and analyzed data. RT, SH, SK, RS wrote and edited the paper. RS supervised research.

## Data availability

The datasets generated and/or analyzed during the current study are available from the corresponding author on reasonable request.

Source data for figures are available from the corresponding author upon request.

