## Supplementary Information for "Endocytosis of ALK promotes glucose uptake in *ALK*-amplified neuroblastoma"

Supplementary Fig 1

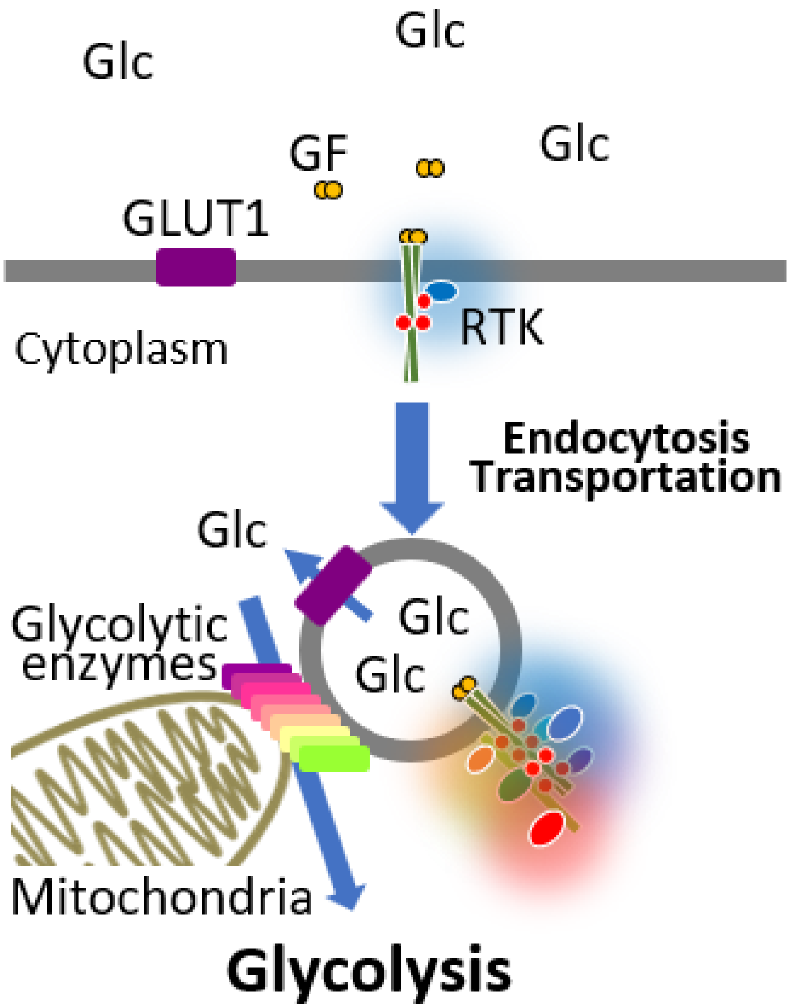


**Fig. S1. RTK endocytosis-dependent glucose uptake.**

Schematic illustrating RTK endocytosis-dependent glucose uptake. GF, growth factor; Glc, glucose; RTK, receptor tyrosine kinase.
